# Quantitative profiling of microbial communities by *de novo* metaproteomics

**DOI:** 10.1101/2020.08.16.252924

**Authors:** Hugo B. C. Kleikamp, Mario Pronk, Claudia Tugui, Leonor Guedes da Silva, Ben Abbas, Yue Mei Lin, Mark C.M. van Loosdrecht, Martin Pabst

## Abstract

Metaproteomics has emerged as one of the most promising approaches for determining the composition and metabolic functions of complete microbial communities. Conventional metaproteomics approaches however, rely on the construction of protein sequence databases and efficient peptide-spectrum matching algorithms. Thereby, very large sequence databases impact on computational efforts and sensitivity. More recently, advanced *de novo* sequencing strategies—which annotate peptide sequences without the requirement for a database—have become (again) increasingly proposed for proteomics applications. Such approaches would vastly expand many metaproteomics applications by enabling rapid community profiling and by capturing unsequenced community members, which otherwise remain inaccessible for further interpretation. Nevertheless, because of the lack of efficient pipelines and validation procedures, those strategies have only rarely been employed for community proteomics.

Here we report on a newly established de novo metaproteomics pipeline which was evaluated for its quantitative performance using synthetic and natural communities. Additionally, we introduce a novel validation strategy and investigate the actual content of community members within community proteomics data.

## INTRODUCTION

State-of-the-art approaches for analysing the composition of microbial communities are based on in-situ staining, 16S ribosomal RNA sequencing or whole-genome shotgun-based approaches. Moreover, metatranscriptomics provides additional gene activity information, but unfortunately, mRNA levels often only poorly correlate with actual protein abundances.^1^ Therefore, above mentioned approaches do not provide insights into the actual phenotype of a community, and the actively expressed pathways responsible for metabolic conversions remain elusive.^2^

Metaproteomics on the other hand, targets the functional parts—the proteins—of a community directly and therefore provides insights into the community phenotype. Furthermore, because proteins make up the bulk mass of a cell, metaproteomics also estimates the contribution of individual community members to the community biomass.^3^

In recent years, metaproteomics has gained substantial momentum with the development of high-resolution proteomics workstations and the establishment of next-generation sequencing (NGS) technologies, which provide affordable high-quality (protein) sequence databases from complete communities.^4^ Classical metaproteomics approaches employ peptide-spectrum matching algorithms used for subsequent protein and species identification. The quality and completeness of employed databases is therefore of utmost importance.^5, 6^ A complete database covers the genetic potential of all community members and therefore may contain hundreds of thousands of sequences. Alternatively, comprehensive (and even larger) public sequence databases such as NCBI, UniProtKB/Swiss-Prot or GenBank may be accessed (in addition),^6^ which however, require advanced focusing/filtering strategies to manage computational efforts.^7-10^ Very large protein sequence databases challenge the common ‘peptide-spectrum matching’ algorithms and associated statistical parameters, which have been historically developed for single-species proteomics. Consequently, conventional metaproteomics experiments can be compromised in regard to sensitivity, accuracy and throughput.^5, 7, 8^

A database-independent approach, which directly annotates mass spectrometric fragmentation spectra with amino acid sequences, such as by *de novo* peptide sequencing, overcomes the above mentioned database-related limitations, and therefore would increase the spectrum of metaproteomics applications.^9^ Advantageously, *de novo* established peptide sequences require only to retrieve taxonomic and functional annotations from comprehensive taxonomic databases using efficient ‘text-search’ functions. Furthermore, *de novo* sequencing avoids loss of taxonomic and functional information from community members not covered by the database. Those signals (not covered by the target database) can be further matched to related species through sequence homology search approaches.^11^ Homology search further increases proteome coverage, by annotating also ‘partially correct’ sequences (sequence tags), which are common ‘by-products’ of the *de novo* sequencing process.^11^

Moreover, *de novo* sequencing may serve as a direct measure of the proportion of unsequenced members in a community. In a similar manner, the usefulness of *de novo* sequencing for evaluating the target sequence database completeness, or ‘suitability’, has been demonstrated only recently.^12^ On the other hand, *de novo* peptide sequencing strongly depends on high-quality mass spectrometric data and efficient sequence annotation tools. Therefore, *de novo* sequencing commonly provides fewer spectral identifications when compared to database search approaches.^13^ Nevertheless, whether *de novo* sequencing provides sufficient qualitative and quantitative information for (quantitative) metaproteomic applications has not been effectively established to date.

Over the past years, several high-performance *de novo* sequencing algorithms have been introduced,^14, 15 16^ and some have also been proposed for taxonomic profiling applications.^17, 18^ In addition, a number of advanced web-based services that support taxonomic and functional analysis of metaproteomic protein and peptide sequences have been introduced only recently. ^19-21, 22, 23^

In this study, we evaluate a newly established *de novo* metaproteomics workflow for its quantitative performance and taxonomic resolution using synthetic and environmental community data. Furthermore, we introduce a validation strategy and demonstrate how to establish the actual content of individual community members within community proteomics data.

## MATERIALS AND METHODS

Proteomics and metaproteomics data: The synthetic community proteomic raw data were downloaded from ProteomXchange server project PXD006118, established by M. Kleiner and M. Strous labs^3^. Protein content and taxonomic lineages of the synthetic community samples used have been further outlined in the supplemental table, sheets 2 and 3. Due to incomplete coverage of viral strains in the Unipept database, viruses were not further considered in the quantitative analysis. Shotgun proteomic raw data from *Rhodopseudomonas palustris* were retrieved from the project PXD013729 generated by E. Nakayasu, Pacific Northwest National Laboratory and C.S. Harwood, University of Washington, *Campylobacter jejuni* raw data were retrieved from PXD005306 generated by M. Monroe, and J. Adkins, Pacific Northwest National Laboratory, Paracoccus denitrificans raw data were downloaded from project PXD013274 generated by T. J. Erb and M. Glatter, MPI Marburg, respectively. *Lactobacillus sakei* PXD011417 from C. Ludwig, Bavarian Center for Biomolecular Mass Spectrometry (BayBioMS), Technical University Munich. Acinetobacter baumannii PXD011302 from M. Feldmann Washington University School of Medicine and J. Scott, University of Melbourne; Streptococcus mutans PXD006735 from J. Koh and K.C. Rice, University of Florida; Trypanosoma brucei PXD009073 from J.W. Dupuy form Centre de Génomique Fonctionnelle Bordeaux, France and M. Boshart, from Ludwig-Maximilians-University Munich Martinsried, Germany. Additional environmental community reference dataset shown in SI Figure 4, was obtained from PXD008780, as published by B. L. Nunn and E. Timmins-Schiffman of the University of Washington^24^.

Peptide sequencing procedures: Mass spectrometric raw data were processed using PEAKS Studio X (Bioinformatics Solutions Inc., Canada)^15^ for database search and *de novo* sequencing, or DeepNovo^14^ for comparative *de novo* sequencing studies. Both, *de novo* sequencing and database search was performed allowing 15ppm parent ion and 0.015Da fragment mass error (depending on the acquisition, slightly more tolerant parameters such as 20ppm/0.02Da were applied). Carbamidomethylation was set as fixed and methionine oxidation as variable modifications. Database search allowed in addition N/Q deamidation as variable modifications. The same settings were applied to DeepNovo where applicable, otherwise software default settings were used. Database search further used decoy fusion for estimation of false discovery rates (FDR) and subsequent filtering of peptide spectrum matches for 1% FDR. Only the top ranked *de novo* sequence annotations were considered for processing.

*De novo* metaproteomics data processing workflow ‘NovoBridge’: A MATLAB (The MathWorks, Inc., US) ‘main script’ was created which links together functions such as pre-filtering, sequence randomisation, collection of taxonomic and functional information, threshold filtering, grouping and data visualisation. Briefly – after top-level quality pre-filtering and additional randomisation of candidate sequences, taxonomic and functional information were collected by programmed submission to Unipept API using ‘pept2lca’ and ‘pept2funct’ commands, equating amino acids I and L. In either case, retrieved taxonomic branches were threshold filtered, grouped and quantitatively accessed using spectral sequence counts or intensities, respectively. Only sequence candidates with an ALC greater than 70 (or for DeepNovo, a *p* score of less than −0.1) and less than 15 ppm mass error were considered for processing. To avoid interference from random assignments, every taxonomic identifier (taxonomic branch) required a minimum of 3 unique identifications before being considered for quantitative assessment. A detailed description of the *de novo* metaproteomics pipeline, including parameters used for functional profiling are outlined in the supplemental information materials. Optionally, peptide sequence lists from high quality unmatched sequences (top 80% based on ALC scores) were processed against the NCBI non-redundant protein sequence database, employing a local installation of BLASTp+ (Basic Local Alignment Search Tool) and the PAM30 scoring matrix. For BLASTp+ searches, the top 5 ranked assignments per query sequence (based on bit score) were collected and further filtered for best e values and score(s).

Reference proteome *in silico* study: A random selection of 3.5K unique trypsin cleaved *in silico* peptides (7-15 amino acids length, to approximate real samples) was generated for every reference proteome listed in the SI table, sheet 4 using MATLAB’s bioinformatics toolbox. The *in silico* peptidomes were processed through the same pipeline as described above.

## RESULTS

The presented metaproteomics pipeline employs conventional high-resolution shotgun proteomics data in which fragmentation spectra are subsequently translated into peptide sequence lists by *de novo* sequencing. The lists are then submitted by programmed access to the (public) Unipept database to retrieve taxonomic and metabolic information.^22^ Annotations are then processed by the established pipeline, which includes grouping into taxonomic branches and translation of enzyme commission numbers into KEGG pathways. We investigated fundamental aspects and evaluated the performance of the established workflow using synthetic and natural microbial communities.

### Taxonomic resolution

The first question concerns the taxonomic resolution that can be achieved when matching *de novo* peptide sequences against particularly large taxonomy databases to retrieve taxonomic and functional annotations. A large number of peptide sequences is common to several taxa and can therefore only be unique to a certain taxonomic ranking. Hence, the number of unique peptide sequences decreases from higher to lower taxonomic rankings. For example, because of the relatedness between taxa, there will be many more peptide sequences unique only to the phylum level compared to the more distinguished genus or species levels.

For our study, we aimed to retrieve taxonomic information from the Unipept database, which contains processed peptide sequences pre-allocated with taxonomic and functional annotations derived from the NCBI taxonomy database.^25, 26^ The Unipept ranking uses the hierarchical structure of the NCBI taxonomy for which consensus taxa have been determined using the lowest common ancestor approach.^25^ To test the Unipept database for the achievable taxonomic resolution, we generated *in silico* peptide sequences from >1000 species retrieved from the NCBI reference sequences database (www.ncbi.nlm.nih.gov/refseq/). This provided for approx. 90% of all peptide sequences taxonomic annotations, but as expected, showed a steady decrease in the number of assigned peptides from higher to lower taxonomic rankings (=‘drop-off rate’), with a particularly large drop between genus and species level (Figure 1C). Noteworthy, deviations from this ‘drop-off rate’ can be observed for species from highly sampled taxa and species with inconsistent taxonomic classifications. This impacts not only on the quantitative performance, but may also limit the taxonomic resolution, because a certain number of peptides is required for the identification of a respective taxon.

**Figure 1.**
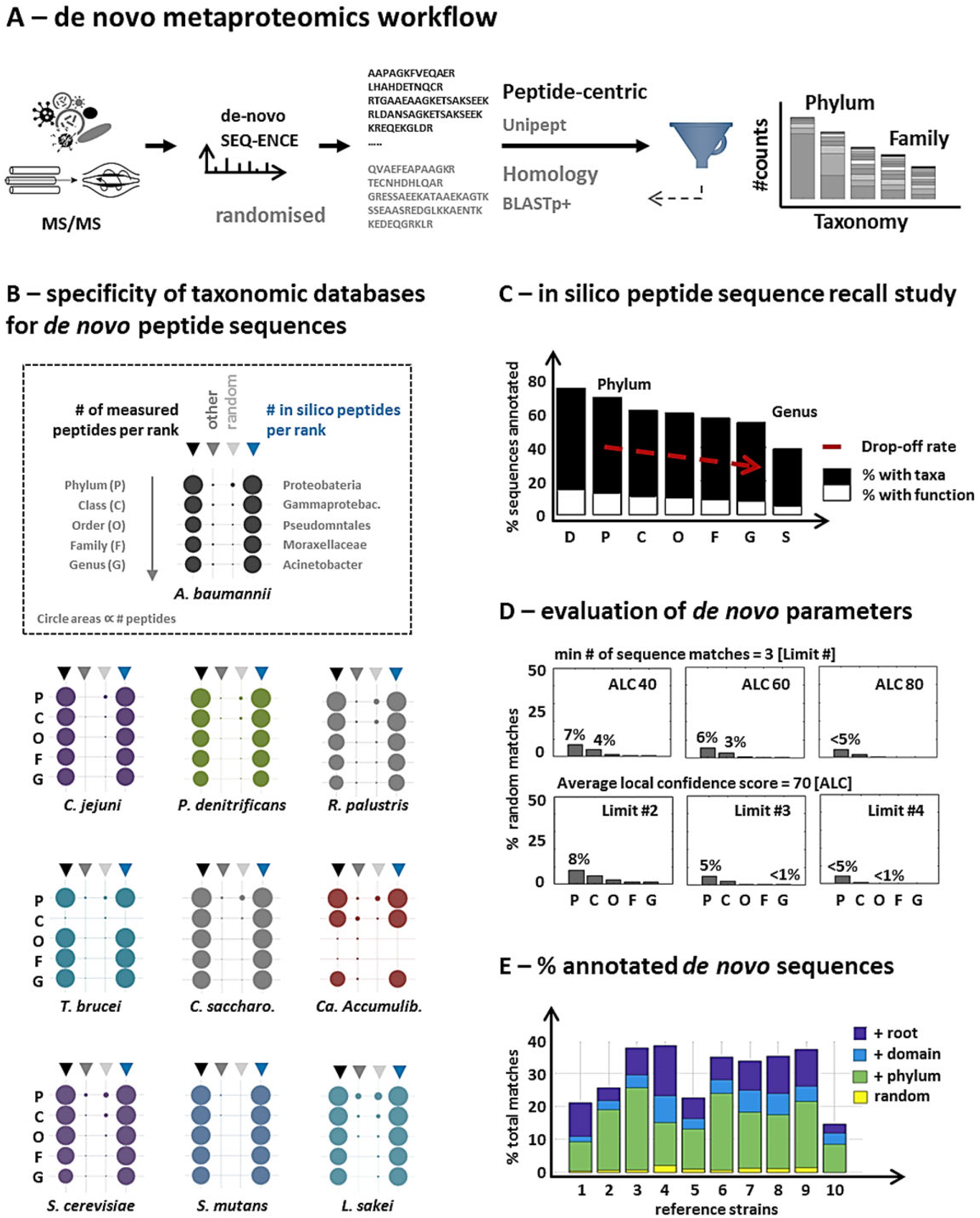
**A) Shotgun metaproteomics workflow.** Shotgun metaproteomic raw data from microbial communities are *de novo* sequenced and processed through the established pipeline as ‘correct’ and randomised sequences. The peptide-centric approach accesses Unipept^26^ to obtain taxonomic and functional annotations. Further processing includes grouping into taxonomic branches and translation of functional annotations into KEGG pathways. High quality unmatched sequences are further made accessible for homology search approaches such as BLASTp+. **B) Specificity of taxonomy databases for *de novo* peptide sequence lists**. Shotgun proteomic data from pure reference strains were *de novo* sequenced and processed through the established *de novo* metaproteomics pipeline to retrieve taxonomic annotations. The annotated sequences were then grouped into taxonomic lineages (phylum – class – order – family and genus) and represented as circle graphs. The circle areas correlate to the normalised sequence counts of the respective taxonomic rank. Every reference strain is represented by four circle lanes: BLACK TRIANGLE ARROW = ‘# of measured peptides per rank’. This counts the number of peptide sequences annotated to the lineage of the target strain, e.g. A. baumannii. GRAY TRIANGLE ARROW: ‘other’. This counts the number of peptide sequences annotated to other taxonomic lineages than the target strain. LIGTH GRAY TRIANGLE ARROW ‘random’. This counts the number of randomised peptide sequences which received a taxonomic annotation. BLUE TRIANGLE ARROW = ‘# of *in silico* peptides per rank’. This counts the number of *in silico* target strain sequences for every rank. The experiment confirms that erroneous or only partially correct *de novo* sequences only insignificantly interfere with the taxonomic representation of the metaproteomic sample. Furthermore, the low number of ‘other’ strain assignments confirmed the purity of the selected reference strain samples. Except for the *in silico* experiments, the averages of duplicate analyses are shown. **C) *In silico* proteome recall study**. The bar graph shows the average number of *in silico* peptide sequences which retrieved taxonomic or ‘enzyme commission number’ annotations. The *in silico* peptide sequences were generated from a large number of proteomes (>1000, retrieved from the NCBI reference proteome database). The individual taxonomic rankings domain (D), phylum (P), class (C), order (O), family (F) genus (G) and species (S) are shown as separate bars. Approx. 90% of the peptides obtained taxonomic annotations (black bars), and 10-20% retrieved additional functional annotations (enzyme commission numbers, white bars). The number of sequence annotations per taxon showed a steady decrease from the phylum to the genus level (‘drop-off’ rate, red arrow). **D) Evaluation of de novo sequence quality parameters**. The bar graph shows the average number random sequences which obtained a taxonomic annotation, when considering different quality parameter thresholds. The randomised sequences were generated from the ‘correct’ reference strains *de novo* sequence lists (excluding *T. brucei* and *Ca*. Accumulibacter). The quality parameter thresholds evaluated were the average local confidence score (ALC, PEAKS platform) and frequency limits (# of peptide sequences observed for an individual taxonomic identifier). ALCs below 60 and frequency limits <3 increased the percentage of random sequence annotations to >5%. Therefore, an ALC of 70 and a minimum of 3 sequence annotations per taxon were set as default thresholds for the experiments in this study. **E) % of annotated de novo sequences**. The bar graph outlines the % of *de novo* sequences submitted to Unipept which retrieved taxonomic annotations. The bars (1 - 10) represent the strains shown in Figure 1B (*A. baumannii*, top of the image - *L. sakei*, bottom right of the image). The blue bars represent all annotations including ‘root’ level – which are sequences common to all domains of life – the light blue bars represent annotations assigned to domain level and lower, and the green bars show annotations assigned to phylum level and lower. The yellow bars indicate the average number of random annotations at the lower taxonomic rankings.

Furthermore, because there is no complete taxonomy database available, there is always a high likelihood of ‘unsequenced’ community members—those that are not in the taxonomy database—being present in the community. Those retrieve annotations through related species mostly at higher taxonomic rankings and will therefore provide only a comparatively low taxonomic resolution.

A quantitative analysis should therefore aim to investigate the ‘drop-off rates’ for individual taxonomic branches, in order to flag poorly quantitative traits. For this, *in silico* peptidomes may serve as highly useful comparators to establish the actual content of a member within the community proteomics data.

### A validation procedure

*De novo* sequencing commonly generates a fraction of only partially correct peptide sequences. This raises the question of whether those incomplete sequences lead to false positive assignments which bias the taxonomic representation of the community.

As a measure of confidence for *de novo* established peptide sequences, the software platform PEAKS provides the average local confidence (ALC) score, and DeepNovo, the p score.^14, 15^ Although these parameters are useful for ranking *de novo* sequences based on their quality, an estimate on the actual number of incorrect sequences is not provided.

Consequently, additional measures are required to give confidence in the taxonomic representation achieved by *de novo* generated sequences. A recently proposed solution employs a taxonomic database containing sequences not only in correct, but also in reverse order. This strategy enables to make use of the widely employed target/decoy approach.^18^ However, database volumes are thereby duplicated, and considering single taxonomic points does not allow to investigate quantitative profiles. Therefore, we aimed not to randomise the target database sequences, but the peptide query sequences instead.

To qualify this approach, we processed proteomics data from pure reference species, once in correct order, and once after peptide sequence randomisation. The randomised sequences retrieved a surprisingly large number of taxonomic annotations at the root (>20%) and super kingdom levels (>10%), but were consistently low for the lower taxonomic rankings (Figure 1B/D). Only small proportions of other taxa were observed, mostly related to culturing and sample preparation conditions, or the samples themselves (such as *virus L-A related proteins* for the yeast *S. cerevisiae)*. Several of those unexpected matches were only identified at certain taxonomic levels, which underlines the importance of measuring complete taxonomic branches rather than single taxonomic points. (Figure 1B, and SI table) Next, we constructed the theoretical drop-off rates using the reference proteomes of the test strains to investigate for ‘hidden’ side populations, not covered by the taxonomic database. This however showed, that the theoretical and the observed drop-off rates were very comparable, which confirmed the purity of the selected reference strains.

In summary, using the pure reference strain samples and the sequence randomisation strategy, we could demonstrate that de novo sequence lists provide only small numbers of erroneous assignments at lower taxonomic rankings (phylum – genus).

### Quantitative community profiling

Finally, we investigated the quantitative aspect when measuring more complex communities. Kleiner et al. (2017) only recently demonstrated the usefulness of metaproteomics for estimating species biomass contributions. Thereby, the authors generated highly useful metaproteomic reference data from synthetic communities, consisting of species with ‘equal protein’ and ‘equal cell’ content. We *de novo* sequenced the publicly available raw data from both synthetic communities, and subjected the obtained sequence lists to our data-processing pipeline. By employing the abovementioned multi-point taxonomic evaluation, we achieved a particularly good quantitative representation of the community as shown for the ‘equal protein’ community (phylum – family) in Figure 2A. The 17 genus level identifiers provided a comparably good correlation, albeit that 3 strains did not provide sufficient unique peptides at this lower level. The same good species abundance correlation was achieved when analysing another dataset of the same ‘equal cell’ community, thereby also comparing 2 different *de novo* sequencing platforms, PEAKS and DeepNovo (Figure 3). Verification of parameters such as ALC scores and mass error, including species abundance correlations obtained for the ‘equal cell’ synthetic community are shown in the SI Figures 1-4.

**Figure 2.**
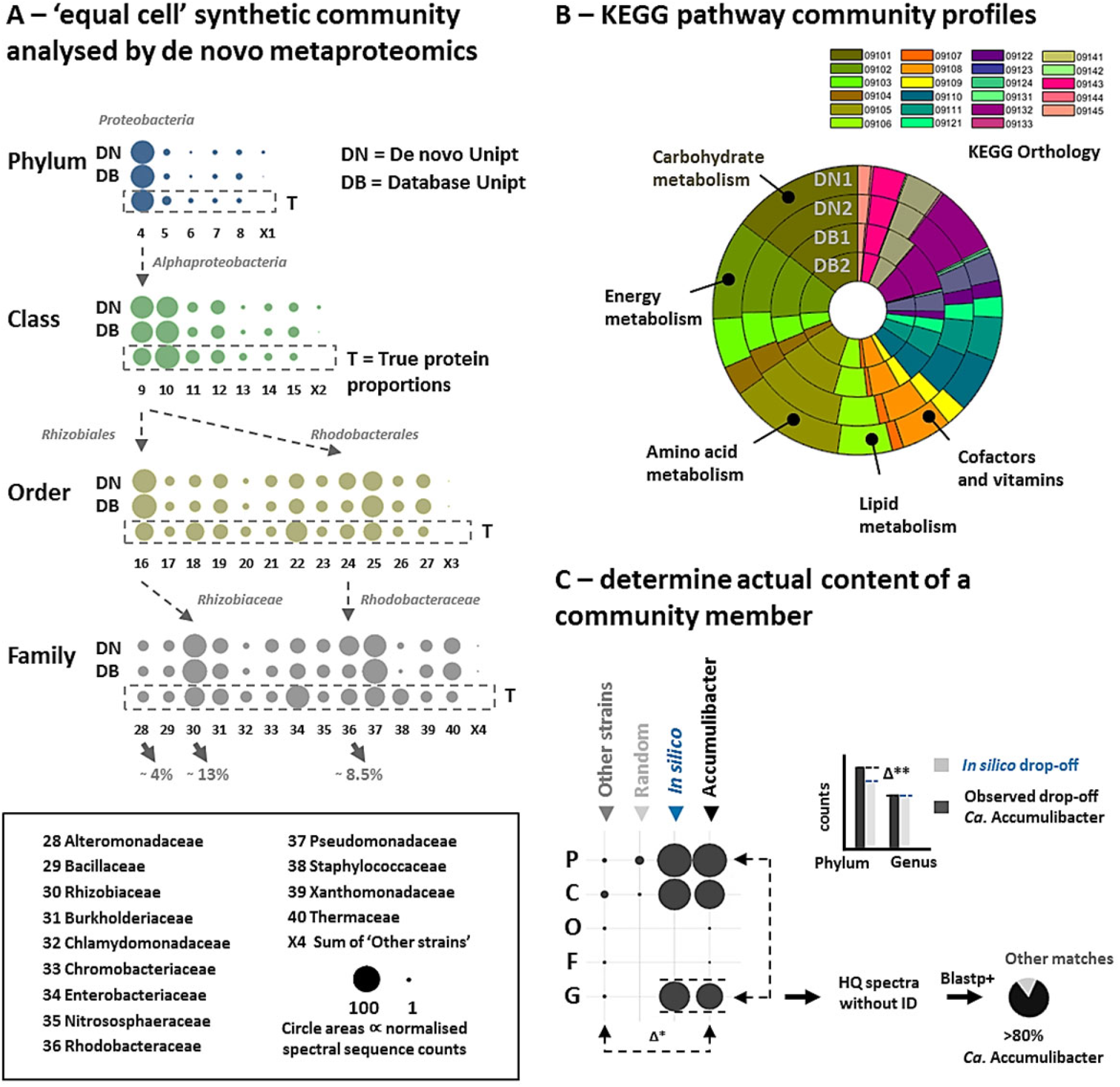
**A) Analysing the community composition by de novo metaproteomics.** Proteomics data from a synthetic community, as established by Kleiner et al., (2017)^3^, were used to evaluate the quantitativeness of the established *de novo* metaproteomics workflow. For this, the raw data were once *de novo* sequenced and once analysed using the constructed target database published by the authors. The taxonomic rankings from phylum – family are represented as circle graphs. Thereby, rows annotated with: ‘DN’ show the protein abundances of each taxon using the *de novo* sequences; ‘DB’ show the protein abundances obtained for each taxon using the sequences established by database matching; ‘T’ show the theoretical (true) protein abundances for each taxon. The circle areas correlate to the normalised spectral sequence counts of the respective taxon. All community members show abundance profiles which strongly correlate to the expected/true (T) species protein abundances. The taxonomic lineages of *Rhizobiaceae* and *Rhodobacteriaceae* are outlined with arrows for exemplification purpose. Those account for approx. 13% and 8.5% of the total community protein content, respectively. Shown is the average of duplicate analyses. **B) KEGG pathway community profiles**. The graphs compare profiles for the major KEGG categories ‘metabolism’ and ‘genetic information processing’, obtained by sequence lists from *de novo* (outer circles) or peptide-spectrum matching approaches (inner circles) of the ‘equal protein’ community. Both, *de novo* (DN) and the database (DB) sequences provide very comparable profiles. Nevertheless, since peptide sequence lists are compared against a large genomic space, sequences can be matched to several enzymes or different pathways, which may inflate functional annotations. See also SI Figure 5. **C) Establishing the actual contribution of community members**. The *de novo* metaproteomic analysis of a *Ca*. Accumulibacter enrichment culture suggests a very high enrichment (>95%, ‘other’ versus ‘Accumulibacter’, Δ*). Furthermore, comparing the experimental with the *in silico* ‘drop-off’ rates, shows only a discrepancy of approx. 17% (small bar graphs, Δ**). To investigate for potential ‘hidden’ members not covered by the taxonomic database, the high quality (HQ) unmatched sequences (top 20% fraction based on ALC scores) were analysed using BLASTp+ for homologue sequences. Thereby, more than 80% of the newly retrieved annotations were again assigned to *Ca*. Accumulibacter (small pie chart), confirming the content estimated after drop-off correction. The individual circle graph columns represent: BLACK TRIANGLE ARROW = ‘# of measured peptides per rank’. This counts the peptide sequences annotated to the lineage of *Ca*. Accumulibacter. BLUE TRIANGLE ARROW = ‘# of *in silico* peptides per rank’. This represents the number of *Ca*. Accumulibacter in silico sequences per taxon. LIGTH GRAY TRIANGLE ARROW ‘random’. This counts the number of randomised peptide sequences which received a taxonomic annotation. GRAY TRIANGLE ARROW: ‘other’. This counts the number of measured peptide sequences annotated to other taxonomic lineages than *Ca*. Accumulibacter. The circle areas correspond to spectral sequence (peptide) counts for the respective taxonomic ranking.

**Figure 3.**
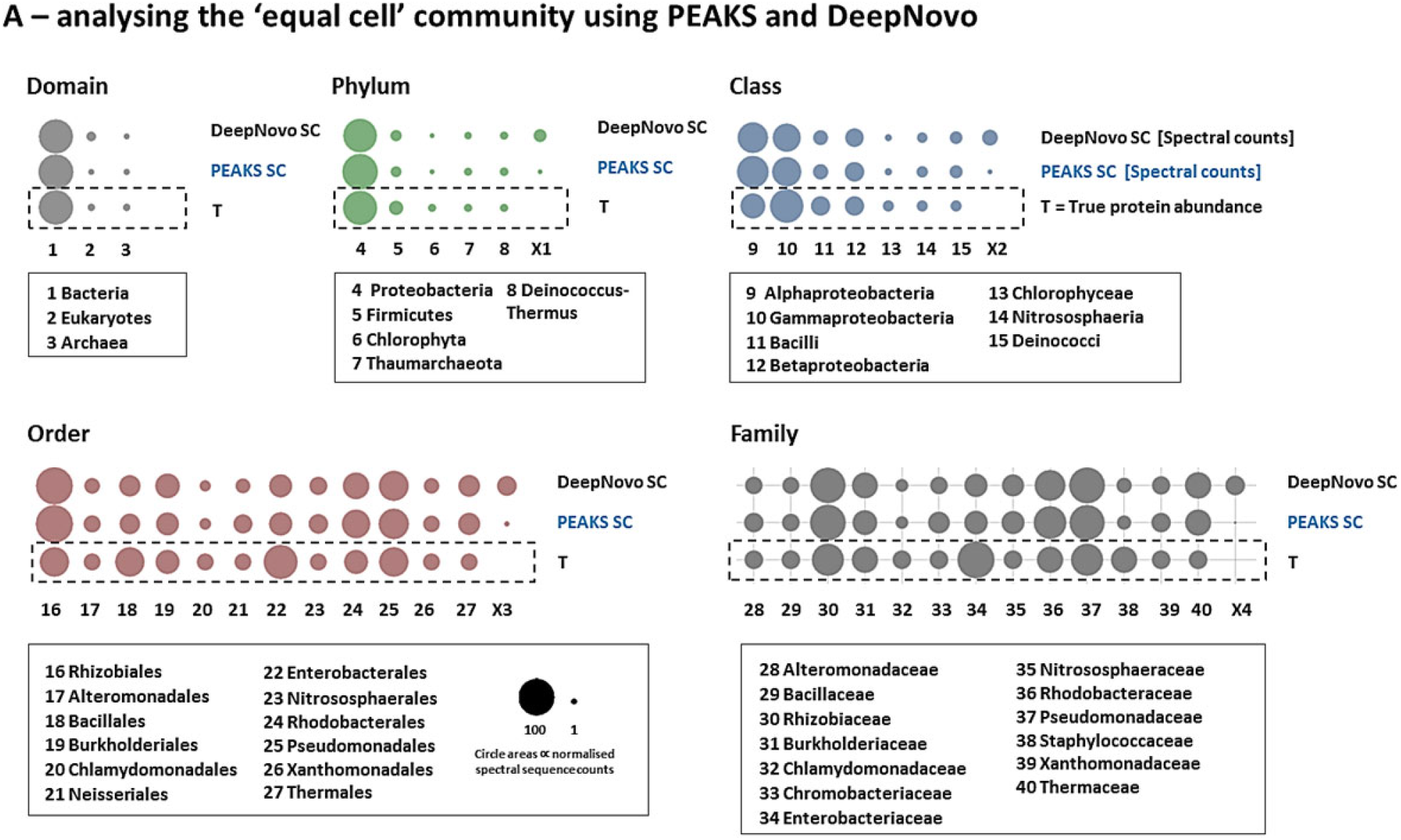
**A) Community profiles of the ‘equal protein’ community established by PEAKS and DeepNovo.** The circle graphs show the taxonomic profiles obtained from the ‘equal protein’ community (Kleiner et al., 2017) established by PEAKS or DeepNovo. *De novo* sequence lists from both platforms were processed by the established de-novo metaproteomics pipeline using the same parameters. ‘T’ represents the true abundance of the respective community members (dashed box). ‘PEAKS SC’ represents the established profiles obtained from the PEAKS *de novo* sequences using spectral sequence counting. ‘DeepNovo SC’ represents profiles obtained from the DeepNovo *de novo* sequences using spectral sequence counting. The unexpected, ‘other’ taxonomic annotations were summed and are shown as circles labelled with ‘X’. The experiment demonstrates that both tools provide very comparable taxonomic profiles and only differ in the proportions of the unexpected ‘other’ matches. The circles represent the average of 2 analyses, where the circle areas correlate to the normalised spectral sequence counts.

Furthermore, we aimed to apply the *de novo* pipeline to a dataset from a natural community. Thereby, we processed a publicly available metaproteomic dataset published by Mikan et al. (2019), representing microbiomes sampled from the Bering Sea.^24^ We generated peptide sequences once using de novo sequencing and once using peptide-spectrum matching employing the metagenomics constructed database published by the authors. Thereby, the taxonomic profiles between both approaches were highly comparable (SI Figure 4), where only some of the very low abundant members were not resolved by the *de novo* approach.

### Establishing the actual content of a community member

Finally, we aimed to investigate the usefulness of in silico drop-off rates and BLASTp+ homology search, for investigating the actual content of an enrichment culture. *Ca*. Accumulibacter has been described frequently as showing strong discrepancies in the proposed community contribution when comparing between FISH- and 16S RNA sequencing-based techniques.^27^ Therefore, we analysed an Accumulibacter enrichment culture metaproteomic dataset through the described pipeline, and observed a particularly high enrichment (Figure 2C, approx. 98% at the genus level (Δ*), in contrast to 16S RNA data for the same reactor at an earlier time point of approx. 34%^28^). When comparing the experimental drop-off rate for the lineage of *Ca*. Accumulibacter with the in-silico constructed drop-off rate, we observed a discrepancy of only approx. 17% (Δ**), meaning that nearly all sequences assigned to proteobacteria translate to the *Ca*. Accumulibacter genus level annotations. Nevertheless, to exclude significant quantities of potential other populations—not captured by the (Unipept) database—the high-quality unmatched sequences (top 20% based on ALC scores) were analysed using BLASTp+ against the non-redundant NCBI protein sequence database (for the sake of speed using a local installation). Thereby, approx. 83% of newly retrieved (genus level) sequences could be attributed again to *Ca*. Accumulibacter (SI-table, Figure 2D), reflecting the estimated content obtained after drop-off correction.

Determining the fraction of unmatched (high-quality) spectra has already been proposed as indicator for the presence of community members not captured by the database.^3, 12^ The fraction of unmatched high quality spectra however, may considerably depend on the applied analytical procedures. The same was observed for the reference strains used in this study, which raw data were acquired in different laboratories, and showed large variations in their fraction of peptides that obtained taxonomic annotations (Figure 1D). Although this approach appears very promising, it may provide misleading conclusions if not corrected for the individual analytical procedures.

## DISCUSSIONS AND CONCLUSION

Metaproteomics has emerged as one of the most promising post-genomics approaches to study microbial dynamics in nature or in the context of human health, such as for the gut microbiome.^5, 16^ However, common metaproteomics workflows require laborious sequence database construction and rely on high quality databases. Thereby, spectrum-matching algorithms are challenged by very large databases or unsequenced community members not covered by the database. Furthermore, the quantitative aspect is often only poorly supported, despite this aspect being of the utmost importance when investigating community dynamics. Here, we describe a newly established *de novo* metaproteomics workflow, which enables quantitative profiling of microbial communities within very short analysis times. We provide a systematic evaluation of the taxonomic resolution and quantitative performance using reference strains and natural communities. Thereby, we introduce a validation procedure, and demonstrate how to establish the actual content of community members within community proteomics data.

The achievable resolution in *de novo* metaproteomics however depends not only on the taxonomic database, but also on the abundance of the individual community members. Most importantly however, the *de novo* metaproteomics approach generates peptide sequences from all community members, which makes ‘not-in-the-database’ sequences accessible for further interpretation using homology search approaches.

## SUPPLEMENTAL INFORMATION

Electronic supplementary information material is available: additional materials and methods, additional supporting figures 1-5, and supporting raw data tables. The Matlab *de novo* metaproteomics pipeline is freely available upon request.

## Supporting information

SI sample info tables

## ACKNOWLEGDEMENTS

The authors are grateful to valuable discussions with our colleagues from the department of Biotechnology and acknowledge Carol de Ram for support with sample processing, and the SIAM consortium for funding.

## REFERENCES

1. Maier, T., Güell, M. & Serrano, L. Correlation of mRNA and protein in complex biological samples. FEBS letters 583, p 3966–3973 (2009).

2. Martin, F. & Uroz, S. Microbial environmental genomics (MEG). (Springer, 2016).

3. Kleiner, M. et al. Assessing species biomass contributions in microbial communities via metaproteomics. Nature communications 8, 1558 (2017).

4. Wilmes, P. & Bond, P.L. Metaproteomics: studying functional gene expression in microbial ecosystems. Trends in microbiology 14, 92–97 (2006).

5. Timmins-Schiffman, E. et al. Critical decisions in metaproteomics: Achieving high confidence protein annotations in a sea of unknowns. The ISME journal 11, 309 (2017).

6. Xiao, J. et al. Metagenomic taxonomy-guided database-searching strategy for improving metaproteomic analysis. Journal of proteome research 17, 1596–1605 (2018).

7. Heyer, R. et al. Challenges and perspectives of metaproteomic data analysis. Journal of biotechnology 261, 24–36 (2017).

8. Muth, T. et al. Navigating through metaproteomics data: a logbook of database searching. Proteomics 15, 3439–3453 (2015).

9. Muth, T., Renard, B.Y. & Martens, L. Metaproteomic data analysis at a glance: advances in computational microbial community proteomics. Expert review of proteomics 13, 757–769 (2016).

10. Potgieter, M.G. et al. MetaNovo: a probabilistic approach to peptide and polymorphism discovery in complex mass spectrometry datasets. bioRxiv, 605550 (2019).

11. Ma, B. & Johnson, R. De novo sequencing and homology searching. Molecular & cellular proteomics 11, O111. 014902 (2012).

12. Johnson, R. et al. Assessing protein sequence database suitability using de novo sequencing. Molecular & Cellular Proteomics (2019).

13. Medzihradszky, K.F. & Chalkley, R.J. Lessons in de novo peptide sequencing by tandem mass spectrometry. Mass spectrometry reviews 34, 43–63 (2015).

14. Tran, N.H. et al. Deep learning enables de novo peptide sequencing from data-independent-acquisition mass spectrometry. Nature methods 16, 63–66 (2019).

15. Ma, B. et al. PEAKS: powerful software for peptide de novo sequencing by tandem mass spectrometry. Rapid communications in mass spectrometry 17, 2337–2342 (2003).

16. Behsaz, B. et al. De novo peptide sequencing reveals many cyclopeptides in the human gut and other environments. Cell Systems 10, 99–108. e105 (2020).

17. Lee, J.-Y. et al. Proteomics of natural bacterial isolates powered by deep learning-based de novo identification. bioRxiv, 428334 (2018).

18. Mooradian, A.D., van der Post, S., Naegle, K.M. & Held, J.M. ProteoClade: a taxonomic toolkit for multi-species and metaproteomic analysis. bioRxiv, 793455 (2019).

19. Mesuere, B. et al. The Unipept metaproteomics analysis pipeline. Proteomics 15, 1437–1442 (2015).

20. Boekel, J. et al. Multi-omic data analysis using Galaxy. Nature biotechnology 33, 137 (2015).

21. Zhang, X. et al. MetaPro-IQ: a universal metaproteomic approach to studying human and mouse gut microbiota. Microbiome 4, 31 (2016).

22. Gurdeep Singh, R. et al. Unipept 4.0: functional analysis of metaproteome data. Journal of proteome research 18, 606–615 (2018).

23. Riffle, M. et al. MetaGOmics: A Web-Based Tool for Peptide-Centric Functional and Taxonomic Analysis of Metaproteomics Data. Proteomes 6, 2 (2018).

24. Mikan, M.P. et al. Metaproteomics reveal that rapid perturbations in organic matter prioritize functional restructuring over taxonomy in western Arctic Ocean microbiomes. The ISME journal 14, 39–52 (2020).

25. Mesuere, B. et al. Unipept: tryptic peptide-based biodiversity analysis of metaproteome samples. vJournal of proteome research 11, 5773–5780 (2012).

26. Mesuere, B. et al. Unipept web services for metaproteomics analysis. Bioinformatics 32, 1746–1748 (2016).

27. Stokholm-Bjerregaard, M. et al. A critical assessment of the microorganisms proposed to be important to enhanced biological phosphorus removal in full-scale wastewater treatment systems. Frontiers in microbiology 8, 718 (2017).

28. da Silva, L.G. et al. Revealing metabolic flexibility of Candidatus Accumulibacter phosphatis through redox cofactor analysis and metabolic network modeling. bioRxiv, 458331 (2018).

