## Supplementary material for "Quantitative profiling of microbial communities by *de novo* metaproteomics": SI text and figures

**Table of content**

1. **Additional materials and methods**
2. **Evaluation of different protein quantification strategies**
3. **Influence of *de novo* sequence quality parameters on taxonomic profiles**
4. **Community profiles of 'equal cell' community**
5. **Bering sea microbiome**
6. **Global community functions using KEGG pathways**
7. **Additional references**

### 1. Additional materials and methods

**Large-scale *in silico* study.** Reference proteomes (>1500) covering all 3 domains of life were retrieved from the NCBI reference database (www.ncbi.nlm.nih.gov/refseq/). *In silico* trypsin cleavage, random selection of 1K sequences (each) and programmed submission to Unipept was done using Matlab2017b (The MathWorks, Inc., US).

**Whole cell lysate proteolytic digestion.** Approximately 25-50mg biomass (wet weight) of each cell pellet/material were homogenised by beads beating in TEAB/B-PER reagent (Thermo Scientific™, for bacterial cells such as *Ca.* Accumulibactor phosphatis enrichment and Clostridium sacch.) or Y-PER reagent (Thermo Scientific™, for yeast cells), respectively. The supernatant was collected by centrifugation at 14.000xg. The protein content was precipitated using TCA (1 vol TCA 100 w/v % to 4 vol sample) followed by washing with ice cold acetone. The protein pellet was resuspended in 200 mM ammonium bicarbonate containing 6M Urea, reduced in a 10 mM DTT solution at 40C for 1 hour, and alkylated using 20 mM IAA in the dark, at room temperature, for 30 minutes. The solution was diluted to below 1 M Urea and digested using sequencing grade Trypsin at a protease to protein ratio of approximately 1:50. Peptides were desalted using Oasis HLB solid phase extraction cartridges (Waters corporation) according to the protocol provided by the manufacturer, speed-vac dried and resuspended in 3% acetonitrile in $H_2O$, containing 0.1% formic acid.

**Shotgun (meta)proteomics.** An aliquot of each sample was analysed using a nano-liquid-chromatography system consisting of an EASY nano LC 1200 equipped with an Acclaim PepMap RSLC RP C18 reverse phase column (50µm x 150mm, 2µm) coupled to a QE plus Orbitrap mass spectrometer (Thermo, Germany). Solvent A was $H_2O$ containing 0.1% formic acid, and solvent B consisted of 80% acetonitrile in $H_2O$, containing 0.1% formic acid. The flow rate was maintained at 300 nL/min. The Orbitrap was operated in top 10 data depended acquisition mode, acquiring peptide signals form 350-1400 m/z, at 70K resolution in MS1 with an AGC target of 3e6 and max IT of 100ms. For yeast, approx. 250ng protein digest were analysed using a short linear gradient from 4 to 30% B over 32.5 minutes, and further to 70% B over 12.5 minutes. MS2 acquisition was performed at 17.5K resolution, with an AGC target of 2e5, and a max IT of 54ms, using a NCE of 28. Unassigned, singly charged as well as 7, 8 and >8 charged mass peaks were excluded. For bacterial samples, approx. 100ng protein digest were analysed using a linear gradient from 5-30% B over 85 minutes and further to 75% B over 25 minutes. MS2 acquisition was performed at 17.5K resolution, with an AGC target of 1e5, and a max IT of 54ms, at a NCE of 30. Unassigned, singly charged, 8 and >8 times charged mass peaks were excluded. Shotgun proteomic raw data have been made available via ProteomeXchange server project PXD016992.

***De novo* metaproteomics pipeline outline (NovoBridge).** A Matlab 'main script' was constructed that links together functions for pre-filtering, sequence randomisation, automated submissions to Unipept to obtain taxonomic and functional information, threshold filtering, taxonomic grouping and visualisation of output data. The pipeline was established and tested with peptide sequence lists generated by *de novo* sequencing using PEAKS or DeepNovo, from high-resolution QE Orbitrap shotgun proteomics raw data. The script was constructed using Matlab 2017b and 2019 respectively. **Function 1, pre-filtering, sequence randomisation and Unipept submission:** The first part of the script involves importing peptide sequence lists (obtained from PEAKS/DeepNovo) into the Matlab environment and to perform pre-filtering based on the sequence annotation quality parameters. The default pre-filtering thresholds were set to ALC scores >40, less than 20ppm mass error and a minimum peptide length of 7 amino acids. Sequence lists were 'cleaned' from peptide modification annotations and mass errors were corrected for mass drifts. The Matlab 'rand' function was further used to generate additional randomised sequences from imported *de novo* lists. Thereby, the order of amino acids in front of the cleavage site (R or K) of every sequence was randomised, keeping original sequence parameters attached. Automated sequence submission to Unipept was done using Unipept`s inbuilt API (https://unipept.ugent.be/apidocs) option.[1] For retrieving taxonomic information, '*pep2lca'* including the options *'&equate_il=true',* to equate leucine and isoleucine, were used. Further, '*&extra=true &names=true'* are specified to get the complete taxonomic lineage and the names of every taxonomic rank. The script automatically filteres for the main categories super kingdom, phylum, class, order family, genus and species. The '*pept2funct'* combined with the option *'&equate_il=true'* was used to retrieve additional EC number information.[1] Thereby, a single peptide sequence can generate multiple EC numbers or pathways which cause functional inference and inflation, particularly when searching against a large sequence database space. For this study, only the top scoring peptide sequence per scan was considered. **Function 2, compositional analysis:** The compositional analysis considered the major taxonomic categories super kingdom, phylum, class, order, family, genus and species. Depending on data quality/abundance, lower ranks (such as species or genus) were excluded from quantitative analysis/representation due to low numbers or insufficient annotations. In a first step, tables were filtered for sequences with ALCs >70 (or less than -0.1 for DeepNovo), and a mass error of less than 15 ppm. To exclude random matches from erroneous *de novo* sequences or low abundant signals, a taxonomic identifier of a branch was only considered when occurring at least 3 times. Frequency and ALC cut-offs/thresholds were established using randomised sequences of the pure reference strains. Remaining taxonomic branches are further grouped and visualised using the 'bar(x..,stacked)' function in Matlab for both, absolute and normalized peptide sequence counts (or areas/intensities, respectively). Visualising the relative abundances of the individual community members were performed using circle graphs using the 'surf' function in Matlab. Circle areas represent thereby the number of normalised spectral sequence counts and show the average of 2 separate analyses (except stated otherwise). True/expected abundances of individual community members of the synthetic communities were retrieved from the supplemental information materials, as published by Kleiner et al., 2017.[2] **Function 3, functional analysis:** KEGG pathways, from global classifications to individual conversions within a pathway, correspond to the KEGG orthology (KO) codes.[3] Therefore, we established a script, which translates the retrieved enzyme commission numbers (EC) into KO codes. This was done by integrating the KEGG annotation database, downloaded from https://www.genome.jp/kegg-bin/get_htext?ko00001 (10/19), into the Matlab environment. The analysis of the global community metabolic functions, considered thereby only branches which were also used for compositional analysis. Sequences assigned to root and super kingdom levels were excluded. EC assignments matched more than twice (based on unique spectral sequence counts) were further translated into KO codes, normalised to the total number of spectral sequence counts and grouped into pathways. Obtained functional community profiles were visualised using heat maps or circle graphs based on KEGG pathways/category levels 2 (global) and 3 (carbohydrate and energy metabolism). Further information regarding 'KEGG pathway categories' are outlined below.[3]*

Heat maps were generated using the 'heatmap' function, and circle graphs were created using Matlab`s 'donut.m' function as available through www.mathworks.com 'file exchange' website.

*Second category codes: 09101 Carbohydrate metabolism, 09102 Energy metabolism, 09103 Lipid metabolism, 09104 Nucleotide metabolism, 09105 Amino acid metabolism, 09106 Metabolism of other amino acids, 09107 Glycan biosynthesis and metabolism, 09108 Metabolism of cofactors and vitamins, 09109 Metabolism of terpenoids and polyketides, 09110 Biosynthesis of other secondary metabolites, 09111 Xenobiotics biodegradation and metabolism, 09121 Transcription 09122 Translation, 09123 Folding, sorting and degradation, 09124 Replication and repair, 09131 Membrane transport, 09132 Signal transduction, 09133 Signalling molecules and interaction, 09141 Transport and catabolism, 09143 Cell growth and death, 09144 Cellular community – eukaryotes, 09145 Cellular community – prokaryotes, 09142 Cell motility.

*Third category codes: 00010 Glycolysis/Gluconeogenesis, 00020 Citrate cycle (TCA cycle), 00030 Pentose phosphate pathway, 00040 Pentose and glucuronate interconversions, 00051 Fructose and mannose metabolism, 00052 Galactose metabolism, 00053 Ascorbate and aldarate metabolism, 00500 Starch and sucrose metabolism, 00520 Amino sugar and nucleotide sugar metabolism, 00620 Pyruvate metabolism, 00630 Glyoxylate and dicarboxylate metabolism, 00640 Propanoate metabolism, 00650 Butanoate metabolism, 00660 C5-Branched dibasic acid metabolism, 00562 Inositol phosphate metabolism, 00190 Oxidative phosphorylation, 00195 Photosynthesis, 00196 Photosynthesis - antenna proteins, 00710 Carbon fixation in photosynthetic organisms, 00720 Carbon fixation pathways in prokaryotes, 00680 Methane metabolism, 00910 Nitrogen metabolism, 00920 Sulfur metabolism. *www.genome.jp/kegg/pathway.html

**Function 4. Peptide sequence outputs.** To interface with other tools, a peptide sequence table output is provided in form of '.xls' or '.mat' files. Thereby either all sequences, only identified or non-identified sequences can be selected. The later can be filtered for high quality spectra, such as selecting for the top 20% (based on ALC score), which was exemplified using the BLASTp+ homology search module, to investigate for potential un-sequenced community members.

***De novo* sequence homology search.** Alternatively, high quality unidentified *de novo* sequences were subjected to BLASTp+ homology search[4, 5]. Even though there are homology search web services available[6], we used a local installation to maintain sufficient throughput and integrity with the established *de novo* metaproteomics pipeline. For this ncbi-blast-2.9.0+ and the non-redundant protein sequence database 'nr.gz' (segmented for more efficient use, due to size) were downloaded from the NCBI ftp server (ftp://ftp.ncbi.nlm.nih.gov/blast, updated 12/19) and installed on a local windows 10 workstation. BLAST searches were operated using the Matlab 'system' command function. All BLAST searches used the PAM30 scoring matrix. Top search results (based on bit-scores) for every sequence were combined and filtered for best e values and scores, respectively. Taxon ID and name databases were downloaded from the NCBI server. Full taxonomic lineages were retrieved form NCBI using E-utilities calls 'http://eutils.ncbi.nlm.nih.gov/entrez/eutils/efetch.fcgi?db=taxonomy&id=' and 'taxurl_right='&retmode=xml'.[7]

### 2. Evaluation of different protein quantification strategies

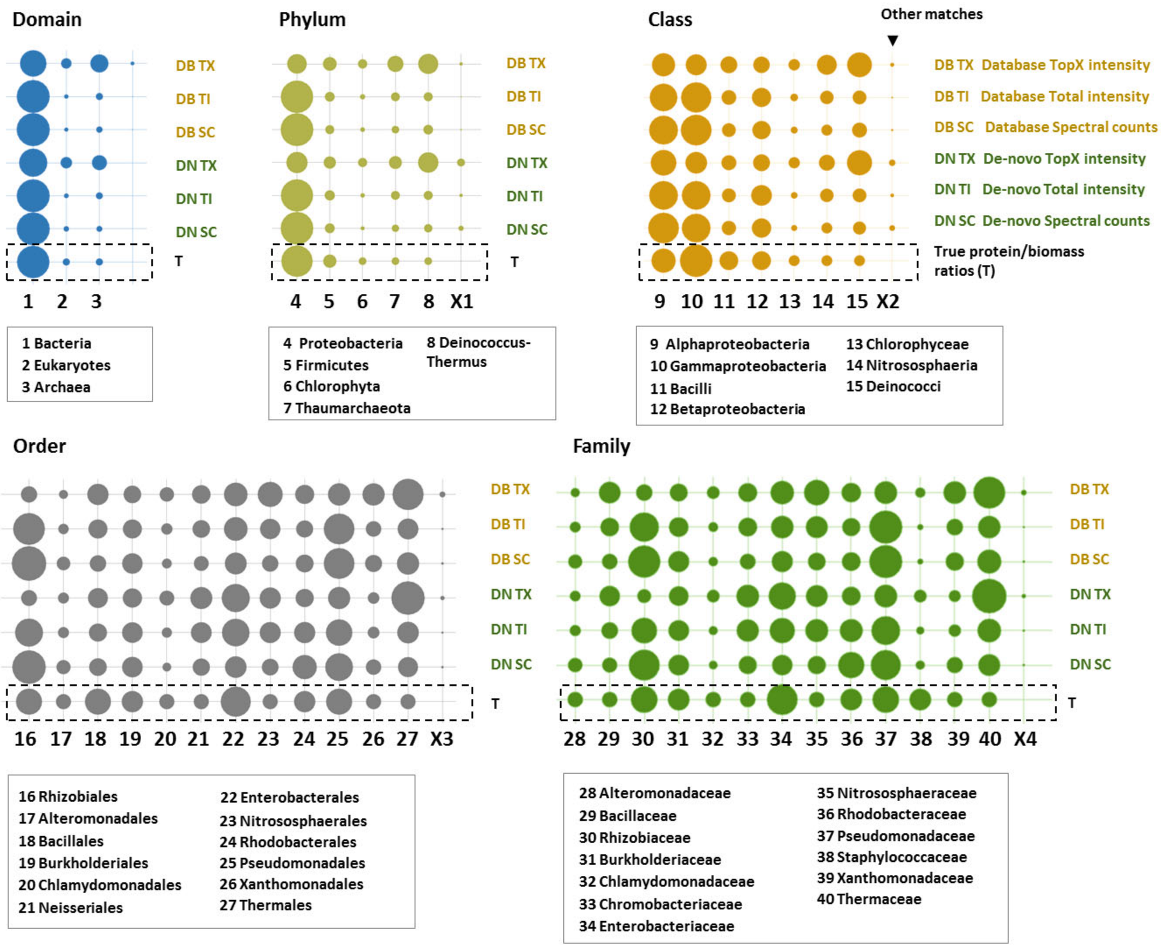


**SI Figure 1:** Taxonomic profiles obtained for the 'equal protein' community (Kleiner et al., 2017), shown from super kingdom to the family level. Quantification was performed using different quantification strategies. Mass spectrometric raw data were processed using *de novo* sequencing or peptide-spectrum matching. Both peptide sequence lists were processed using the established *de novo* metaproteomics pipeline. 'T' represents the true, or expected abundances of the respective community member(s), surrounded by a dashed box. 'DN SC' shows abundances obtained from the *de novo* sequence lists using spectral sequence counting. 'DN TI' shows abundances obtained from the *de novo* sequence lists using the total spectral (peptide) intensities. 'DN TX' shows abundances obtained from the *de novo* sequences using the intensity of the top 5 most intense spectra. 'DB SC' shows abundances obtained from the database search sequences using spectral sequence counting. 'DB TI' shows abundances obtained from the database search sequences using the total sequence (peptide) intensities. 'DB TX' shows abundances obtained from the database search sequences using the intensity of the top 5 most intense spectra.
The sum of 'other' taxonomic annotations is shown as circles annotated by 'X'. Best correlation was observed for the spectral sequence counting strategy, whereas the 'top 5' approach showed the poorest correlation.

#### 3. Influence of de novo sequence quality parameters on taxonomic profiles

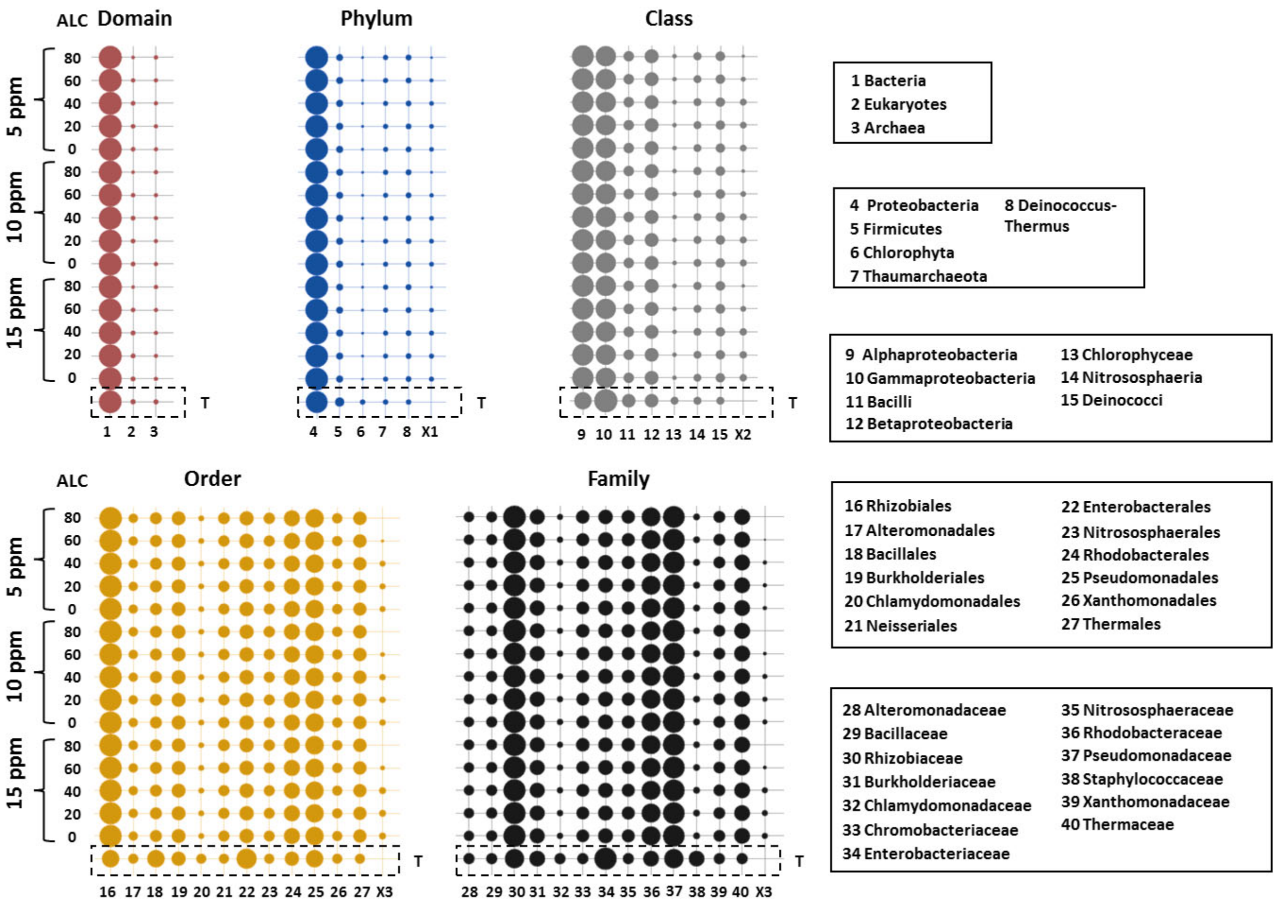


**SI Figure 2:** The influence of the *de novo* sequence quality parameters 'average local confidence score' (ALC) and mass error (Δ ppm, of the proposed sequence) on quantitative taxonomic profiles was investigated using the 'equal protein' community (Kleiner et al., 2017). Mass spectrometric raw data were *de novo* sequenced and processed by the established *de novo* metaproteomics pipeline. The taxonomic profiles were visualised as circle graphs, where the circle areas correlate to the normalised spectral sequence counts. Shown are the average of 2 separate analyses. The sum of unexpected 'other' taxonomic annotations is shown by circles labelled with 'X'.

The experiment demonstrates that the obtained taxonomic profiles were comparatively constant and close to the true abundances ('T') throughout the investigated parameter settings. However, the quantities of 'unexpected' taxonomic annotations decreased when considering only *de novo* sequences with an ALC score >60.

#### 4. Community profiles of 'equal cell' community

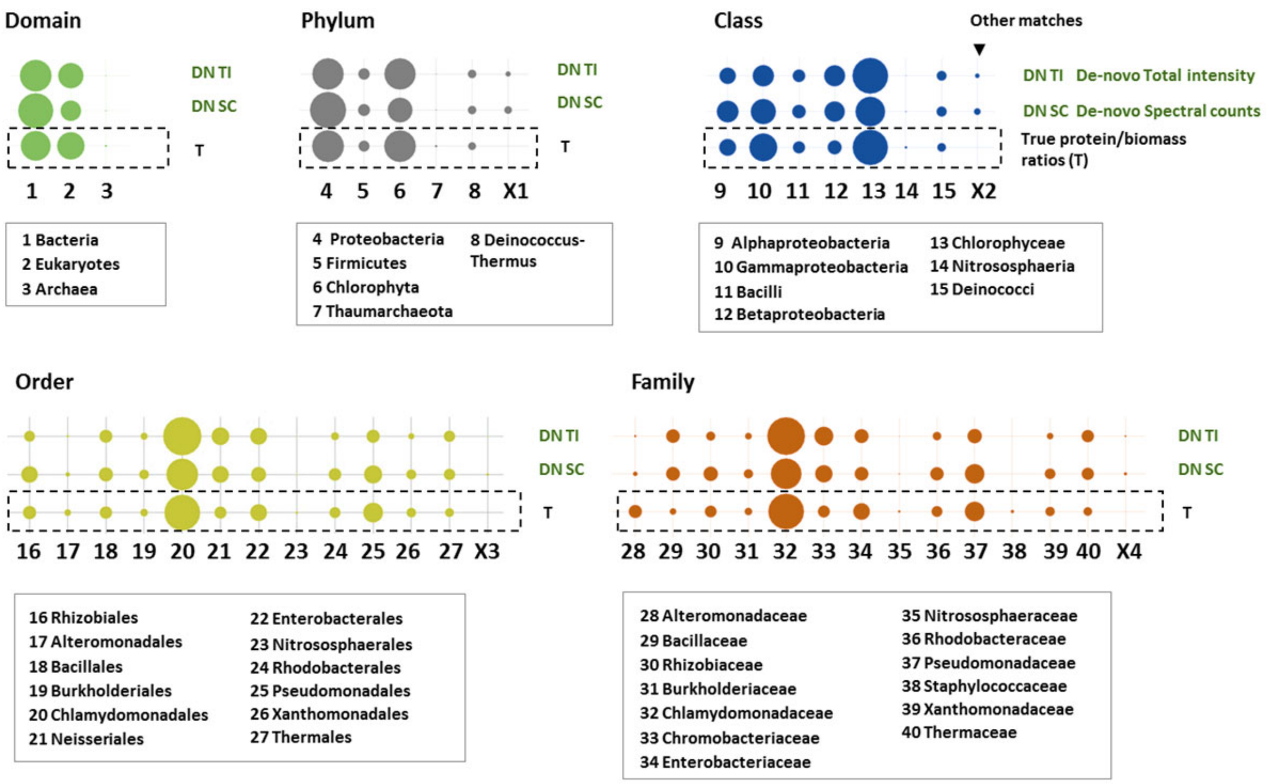


**SI Figure 3**: The above circle graphs show the quantitative profiles of the 'equal cell' community (Kleiner et al., 2017) with the aim to investigate a community with large protein abundance differences. Mass spectrometric raw data were processed by the described *de novo* metaproteomics pipeline, as described in the materials and methods section. The quantification was performed by either summing up spectral sequence counts or by summing up spectral intensities. 'T' represents the true, or expected abundances of the respective community member(s). 'DN SC' shows abundances obtained from the *de novo* sequences using spectral counting (sum of all sequence annotations). 'DN TI' shows abundances obtained from the *de novo* sequences using the total peptide intensity (sum of all sequence intensities). The sum of 'other' taxonomic annotations are shown by circle graphs labelled by 'X'. Overall, the observed community profiles were highly comparable to the true/expected profiles ('T'). Except *Staphylococcaceae* (#38, family level), all other taxonomic identifiers (from phylum - family) could be observed and were close to the true/expected abundance profiles. The circles areas represent the normalised spectral sequence counts of the respective taxonomic identifiers. Shown is the average of 2 separate analyses.

### 5. Bering Sea microbiome

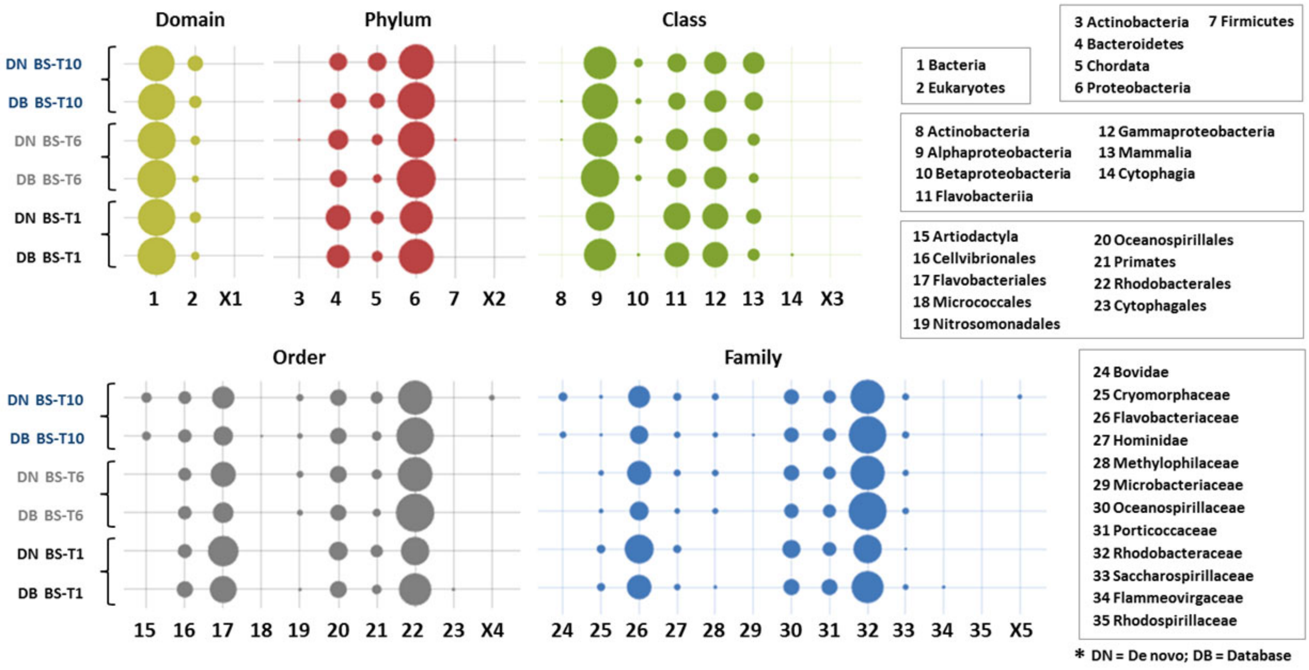


**SI Figure 4:** The circle graphs show the obtained community profiles obtained for the different marine community samples BS-T1, BS-T6 and BS-T10, which were sampled from the Bering sea, as published by Mikan et al (2019).[8] Metaproteomic raw data were retrieved from the proteome exchange server, *de novo* sequenced and analysed by the established *de novo* metaproteomics pipeline. The circle graphs labelled with 'DN' show the community profiles obtained from the *de novo* sequence lists using spectral sequence counting. 'DB' shows the community profiles obtained from sequences of the comparative peptide-spectrum matching experiment, using the protein sequence database published by the authors.

The overall taxonomic profiles obtained from both approaches are highly comparable, where differences are only found for the very low abundant community members, at the lower taxonomic rankings. The circle areas correlate to the normalised spectral sequence counts of the respective taxonomic identifiers. Taxonomic annotations only observed for the *de novo* sequences were summed and are shown as circles annotated by 'X'. The graph shows the average of 2 separate analyses.

#### 6. Global community functions visualised as KEGG pathways

Since GO terms have been reported challenging for enrichment analyses due to unclear hierarchies and dependencies,[9, 10] our workflow translates the obtained EC assignments into KO terms and KEGG pathways. Although the peptide-centric approach compares sequences to a large genomic space (sequences may be matched to several enzymes and different pathways), the overall metabolic profiles appear very comparable between peptide-spectrum matching and *de novo* generated sequence lists. (Figure 2B, and SI Figure 5) Nevertheless, the large database space may limit taxonomic resolution and inflate functional annotations. An investigation with deeper taxonomic resolution requires to maximise proteome coverage and to increase the numbers of unique assignments at lower taxonomic rankings. This is supported through spectrum-matching approaches using tailored databases or potentially also by extensive homology search on high quality sequences. EC numbers were translated into KEGG Orthology (KO) pathways, where only sequences which were kept for taxonomic analysis, were considered for evaluation of the functional profiles. Higher taxonomic rankings, such as annotations to the root or super-kingdom levels were not considered. 2 unique annotations per EC number were required as minimum. Thresholds were generally investigated/tested by using randomised peptide sequences.

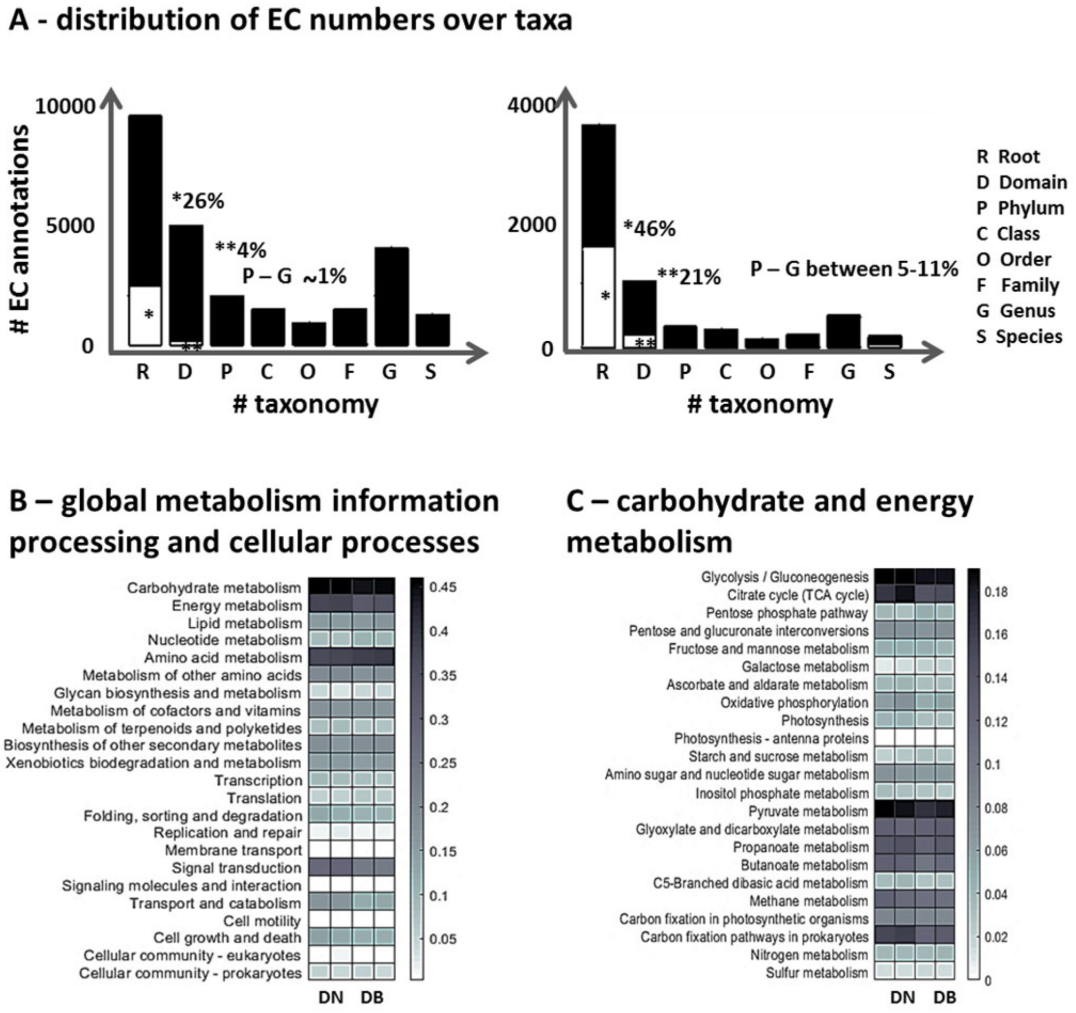


**SI Figure 5. A)** The bar graphs visualise the distribution of enzyme commission number (EC) assignments for database search (A, left bar graph) and *de novo* sequencing (B, right bar graph) generated sequence lists. Annotations from ‘correct’ sequences are shown by black bars and EC annotations retrieved from the randomised sequences are represented by grey bars.

Random EC assignments were frequently retrieved for ‘root’ and ‘super kingdom’ levels. Therefore, EC annotations from those taxonomic levels were excluded from functional analysis. The total number of assignments (phylum to species level) were lower for the *de novo* sequence lists, which is in agreement with the lower number of spectral assignments for *de novo* sequencing, compared to database search approaches. The profiles have been established from the ‘equal protein’ community as provided by Kleiner et al., 2017.[2]

**B/C)** Global functional community profiles of the ‘equal protein’ community, established by sequence lists from database search (DB) and *de novo* (DN) sequencing. EC numbers retrieved from Unipept were translated into KO codes, grouped into KEGG pathways[#] and visualised using heat maps. **A** shows the global profiles for the KEGG ‘categories’ (1) metabolism, (2) genetic information processing, (3) environmental information processing and (4) cellular processes. **B** further details the metabolic profiles for the (1.1) carbohydrate metabolism and the (1.2) energy metabolism.

This experiment shows very comparable profiles between the different peptide sequence annotation approaches. Clustering into KEGG pathways provides a well-structured way to visualise overall community profiles, but may also lead to inflating pathways, because sequences may retrieve more than one EC number or pathway annotation, respectively. The large sequence space of generic databases may further lead to multiple annotations for a single sequence, thereby lowering the accuracy of metabolic profiles.

The above shown community profiles were generated from the 'equal protein' community raw data as established by Kleiner et al., 2017.[2] #https://www.kegg.jp/kegg/pathway.html#metabolism

### 7. Additional references


1. Mesuere, B. et al. The Unipept metaproteomics analysis pipeline. *Proteomics* **15**, 1437-1442 (2015).
2. Kleiner, M. et al. Assessing species biomass contributions in microbial communities via metaproteomics. *Nature communications* **8**, 1558 (2017).
3. Kanehisa, M. & Goto, S. KEGG: kyoto encyclopedia of genes and genomes. *Nucleic acids research* **28**, 27-30 (2000).
4. Madden, T. in The NCBI Handbook [Internet]. 2nd edition (National Center for Biotechnology Information (US), 2013).
5. Christiam Camacho, T.M., Tao Tao, Richa Agarwala, Aleksandr Morgulis BLAST® Command Line Applications User Manual. *Bookshelf NCBI* (2008).
6. Junqueira, M. et al. Protein identification pipeline for the homology-driven proteomics. *Journal of proteomics* **71**, 346-356 (2008).
7. Sayers, E. The E-utilities in-depth: parameters, syntax and more. *Entrez Programming Utilities Help [Internet]* (2009).
8. Mikan, M.P. et al. Metaproteomics reveal that rapid perturbations in organic matter prioritize functional restructuring over taxonomy in western Arctic Ocean microbiomes. *The ISME journal* **14**, 39-52 (2020).
9. Gaudet, P. & Dessimoz, C. in The Gene Ontology Handbook 189-205 (Humana Press, New York, NY, 2017).
10. Simopoulos, C.M. et al. pepFunk, a tool for peptide-centric functional analysis in metaproteomic human gut microbiome studies. *bioRxiv*, 854976 (2019).
